# Emergence of counting in the brains of 3- to 5-year-old children

**DOI:** 10.1101/2022.12.13.520249

**Authors:** Alyssa J. Kersey, Lauren S. Aulet, Jessica F. Cantlon

## Abstract

Learning to map number words onto their ordinal and quantitative meanings is a key step in the acquisition of formal mathematics. Previous neuroimaging work suggests that the intraparietal sulcus (IPS), the inferior frontal gyrus (IFG), and the fronto-temporal language network may be involved in representing number words. However, the contribution of early-developing numerosity representations to the acquisition of counting has not been tested in children. If regions that support numerosity processing are important for the acquisition of counting, then there should be functional overlap between numerosity representations and number word representations in the brain, before children have mastered counting. Using functional magnetic resonance imaging (fMRI), we identified numerosity processing regions in 3- to 5-year-old children during a numerosity comparison task. To identify neural representations of number words, we measured changes in neural amplitudes while those same children listened to number words and color words and while they listened to counting and alphabet sequences. Across multiple whole-brain analyses, we found that the bilateral IPS consistently supported representations of numerosities, number words, and counting sequences. Functional overlap between numerosities and unknown counting sequences was also evident in the left IFG, and in some cases number word representations emerged in the left hemisphere fronto-temporal language network. These results provide new evidence from children that primitive numerosity processing regions of the brain interface with the language network to ground the acquisition of verbal counting.

**Highlights:**

- fMRI data revealed the neural basis of counting acquisition in 3- to 5-year-olds.
- Overlap between neural responses to count words and numerosity emerged in the IPS.
- Sensitivity to number words emerged in the IPS across two different tasks.
- Number word stimuli also engaged regions of the language network in children.
- The IPS and language network may ground number words during counting acquisition.

## 1. Introduction

Preverbal and verbal number concepts develop predictably and tractably in children. Children have robust and abstract nonverbal representations of quantity (numerosities) beginning in infancy (Izard et al., 2009; Lipton & Spelke, 2003), but they do not begin to learn formal verbal counting until around 2.5 years. A central focus of research in numerical cognition concerns what role nonverbal numerosity representations play in the acquisition of symbolic number (Carey et al., 2017; Odic et al., 2015; vanMarle et al., 2018). Theories of cognitive development predict that higher level cognition builds on basic, “core”, concepts (Gallistel, 1990; Spelke & Kinzler, 2007), and theories of concept representation suggest that symbolic concepts are grounded in basic perceptual and motor representations (Barsalou, 1999; Binder & Desai, 2011; Mahon & Caramazza, 2008). In line with these theories, accumulating neural evidence suggests that the cultural acquisition of mathematics, including numerical symbols, builds on primitive numerosity representations in the intraparietal sulcus (IPS) (Ansari, 2008; Cantlon et al., 2006; Dehaene & Cohen, 2007; Nieder & Miller, 2003).

The IPS is a strong neural candidate for counting acquisition because numerosity representations are “native” to that region, as evidenced by neural tuning to numerosities in non-human animals, human adults, and children as young as three years old (Kersey & Cantlon, 2017b; Nieder, 2016; Piazza et al., 2004). In young children who are in the early stages of symbolic number acquisition, the IPS is the primary brain region that shows evidence of numerosity tuning. A causal role for the IPS in numerical and mathematical processing is suggested by patient data showing that damage to IPS impairs calculation (e.g., Dehaene & Cohen, 1997; Takayama et al., 1994), data showing that children with dyscalculia have differential engagement and anatomy of IPS (e.g., Molko et al., 2003; Price et al., 2007), and data showing numerical processing deficits similar to dyscalculia following transcranial magnetic stimulation (TMS) to parietal cortex (Cohen Kadosh et al., 2007).

There is also some evidence that prefrontal cortex is involved in the acquisition of numerical symbols. For one, human and nonhuman primates show evidence of primitive representations of numerosities in this region (Jacob & Nieder, 2009; Viswanathan & Nieder, 2013), though in humans prefrontal regions do not consistently exhibit neural tuning to numerosity (Kersey & Cantlon, 2017b; Piazza et al., 2004). In nonhuman primates who have been trained to recognize numerical symbols, it is primarily prefrontal neurons, and to a lesser degree parietal neurons, that associate visual symbols with numerical values (Diester & Nieder, 2007; Nieder, 2009). In humans, neuroimaging studies reveal that older children who know how to count show stronger recruitment of prefrontal cortex for numerical symbols compared to adults (Ansari et al., 2005; Cantlon et al., 2009a; Rivera et al., 2005), though both the IPS and the prefrontal cortex respond selectively to numerical stimuli in children and adults (Arsalidou et al., 2017; Arsalidou & Taylor, 2011; Kersey & Cantlon, 2017a).

The IPS and prefrontal cortex represent both numerosities and symbolic numbers in proficient counters (Cantlon et al., 2009a; Holloway & Ansari, 2010; Kersey & Cantlon, 2017a; Lussier & Cantlon, 2017; Piazza et al., 2007). However, it is unclear whether children’s early acquisition of number symbols/words involves non-symbolic numerosity representations. One possibility is that early number word semantics are supported by numerosity representations in the IPS and/or prefrontal cortex (Park & Brannon, 2014). If this is the case, then even number words that children have not yet mastered (i.e., numbers outside of children’s counting range), but recognize as numerically relevant should activate regions that represent numerosity. Alternatively, another possibility is that early number word semantics are primarily supported by linguistic representations, and do not involve numerosity representations in IPS and/or prefrontal cortex (Carey & Barner, 2019). If this is the case, then only number words that children have mastered (i.e., numbers within children’s counting range) should activate regions that represent numerosity.

In the current study, we presented spoken number words to 3- to 5-year-old children in two functional magnetic resonance imaging (fMRI) paradigms: a ‘Word Processing’ task and a ‘Natural Counting’ task. As controls, we also presented color words in the ‘Word Processing’ task and alphabet sequences in the ‘Natural Counting’ task. To investigate how the brain supports number word acquisition, we identified neural representations of number words across these two tasks, tested for correlations between children’s neural patterns and behavioral indices of their knowledge, and compared neural representations of number words within children’s counting range (“Known number words”) to neural representations of number words outside children’s counting range (“Unknown number words). Children’s number knowledge was assessed in three ways: by their memorized counting, acquisition of the cardinality principle, and general knowledge of how to use and manipulate numbers (see the Results and the Methods for more details). Regions that are identified consistently across tasks and analyses are likely to be important for the semantic acquisition of number words. Across multiple whole-brain analyses, we evaluated the hypothesis that symbolic number representations build on primitive numerosity representations by testing for functional overlap between neural representations of count words and neural representations of numerosities, identified using a ‘Numerosity Discrimination’ task.

## 2. Methods

### 2.1 Participants

68 typically developing children (3.06 – 5.96 years) and 20 adults (20.2 – 31.6 years, mean age = 23.9 years, 10 women) participated in this study. All participants had normal or corrected to normal vision and no history of neurological impairments. Adult participants and the parents of all children gave informed written consent in accordance with the University of Rochester’s Research Subjects Review Board. Children were excluded from the analyses for failure to complete any functional scans (n = 5), for excessive motion (> 3 mm of volume-to-volume motion or > 5 mm of overall displacement, n = 1 child from Word Processing Task, n = 19 children on one or more runs of the Numerosity Discrimination Task, n = 15 children from Natural Counting), for inattention or noncompliance during experimental tasks (n = 5 from Natural Counting) or for low accuracy on the numerosity discrimination task (accuracy < 60%, n = 1 run from 5 children, n = 2 runs from 8 children). This resulted in a sample of 45 children who successfully completed 1 or 2 runs of numerosity discrimination task (19 girls, mean age = 4.7; n=13 with 1 run, n=32 with 2 runs), 43 children who successfully completed the natural counting task (23 girls, mean age = 4.6), and 16 children who successfully completed the word processing task (12 girls, mean age = 4.7; n = 4 with 1 run, n = 11 with 2 runs, n = 1 with 3 runs; see Table 1). The maximum amount of motion within a run in the data from the included children was less than 3 mm (average motion = 0.33 mm; range of average motion = 0.02– 2.60 mm; see *Preprocessing* for motion correction methods).

**Table 1.** Included data from child participants by task.

|  | **Numerosity**<br>**Discrimination** | **Natural**<br>**Counting** | **Word**<br>**Processing** |
| --- | --- | --- | --- |
| **Usable N** | 45 | 43 | 16 |
| **Girls/Boys** | 19/26 | 23/20 | 12/4 |
| **Mean Age (years)** | 4.7 | 4.6 | 4.7 |
| **Mean Number of**<br>**Runs per Child** | 1.7 | 1 | 1.8 |
| **Total Possible Runs** | 2 | 1 | 3 |

### 2.2 fMRI Session

Prior to scanning, children were familiarized with the MRI environment, practiced lying still in a mock scanner, and practiced the Numerosity Discrimination Task and Word Processing Task. Following the 30-min practice session, children proceeded to the actual MR scanner where their heads were secured with headphones, foam padding, and medical tape. Adults simply received verbal instructions about the tasks.

Stimuli were presented using MATLAB R2014B and the Psychophysics toolbox.

### 2.3 Word Processing Task

To identify regions involved in word processing in early childhood, number words ranging from 1 to 99 and color words were verbally presented in mini-blocks. Stimulus blocks consisted of 4 words from the same category (number, color) presented aurally at a rate of 1 word/2 sec. Words were divided into Known number words, Unknown number words, Known color words, and Unknown color words using children’s behavioral scores. Number words within the child’s counting ranges were categorized as “Known” and number words beyond the child’s counting ranges were categorized as “Unknown”, even though children recognize “Unknown” number words as numerically relevant (Wang & Feigenson, 2019). For example, for a child that could count to 20, the number “thirty-seven” was classified as Unknown, even though the child knew the meaning of “seven” and likely identified “thirty-seven” as numerically relevant. Up to twelve Unknown number words and Known number words were randomly selected for each child.

Color words were selected from sets of “Known” and “Unknown” color words to match the number of syllables in the corresponding number word lists (see Supplement 1, Table S1 for a full list of color word stimuli). To obtain the necessary number of syllables, Known colors words included simple compound color words (e.g., “light pink” and “dark purple”) and words that were a combination of Known colors words (e.g., “bluish-green”). For Unknown color words, 40% of the words were compound color words that consisted of a known color word with an unfamiliar word (e.g., “ash gray”). The remaining color words were complex color words (e.g., “lavender” and “wisteria”). Compound color words were included to mirror Unknown number words, such a child likely will recognize “gray” as a color but will be unsure about the meaning of “ash.” Runs consisted of 3 stimulus-blocks for each stimulus category separated by interblock intervals of 10-seconds. Stimulus-blocks were randomized such that no category was presented back-to-back.

To keep children engaged during listening, children completed an incidental task in which they were instructed to press a button when they heard the word “boat.” The word “boat” was embedded in 3 catch-blocks: the word “boat”, a color word, and a number word were presented in a random order. Number and color words that were presented in these catch-blocks were excluded from analyses. Catch-blocks were evenly distributed throughout the run (1 catch-block every 4-6 blocks). To help children remember what they were listening for, they were told to fixate on a small boat image in the top center of the screen. The boat-fixation appeared on the screen after 12-seconds of a blank screen, was presented without auditory stimuli for 2-seconds, and then remained on the screen during the presentation of the stimuli and during the inter-block intervals. After the 12 stimulus blocks and 3 catch blocks had been presented, the experiment ended with an additional 12-seconds of a blank screen. Total run time was 4.67 min (140 volumes at TR = 2 sec).

### 2.4 Natural Counting Paradigm

In the natural counting paradigm (Supplement 2), child and adult participants were instructed to carefully listen to pre-recorded audio tracks of a person counting or saying the alphabet. The alphabet was used as a comparison to the count list because it represents another ordinal sequence of symbols that children learn to recite around the same age that they learn to recite the count list. Sequences were presented in 70-second blocks of 60 items presented at a rate of 1 item every 1.1 - 1.2 seconds. Fifteen blocks were presented throughout the experimental paradigm and were separated by 4-seconds of blackscreen. To keep the child participants engaged in the natural counting task, short clips from child-friendly cartoons were presented on the screen. Audio tracks were removed from the cartoons and were replaced with quieter, child-friendly instrumental music. Cartoon tracks and music were identically matched across number and alphabet sequences and were played for all participants (including adults). The scan began and ended with 12 seconds of blackscreen resulting in a total scan time of 18.9 minutes (567 volumes at TR = 2 sec).

Counting sequences consisted of three blocks of 1 to 10 repeated 6 times per block, three blocks of 11 to 20 repeated 6 times per block, and three blocks of higher sequences of 30 items repeated twice per block (one block of 51 to 80, one block of 61 to 90, and one block of 71 to 100). Alphabet sequences consisted of three blocks of A to J repeated 6 times per block (10 items per sequence) and three blocks of the entire alphabet repeated twice all the way through and then a third time up to H (26 items per sequence).

### 2.5 Numerosity Discrimination Paradigm

To localize the numerosity processing network, children and adults completed a numerosity discrimination paradigm. Two arrays of cookies were presented on either side of a fixation cross. One array was paired with Cookie Monster, and one array was paired with Oscar the Grouch. Participants were instructed to press a button on the same side as whoever had the most cookies^1^. Comparisons were made across an easy ratio (0.25) or a difficult ratio (0.60). During the mock scanner practice session, all comparisons were made across an even easier ratio (0.20). One run consisted of 48 trials divided into 16 mini-blocks (3 trials per mini-block). Trial order was varied across participants. Arrays were presented for 2 seconds with a 2-second inter-trial interval and a 6-second inter-block interval. The experiment began with 12-seconds of blackscreen, followed by 2-seconds of a fixation cross and ended with another 12-seconds of blackscreen. Total time for one run was 4.6 minutes (139 volumes at TR = 2 sec).

Cookie arrays were presented on a black background in the top center of the viewing screen to the left and the right of a central fixation cross. Arrays consisted of 1 to 30 cookies. Oscar the Grouch was always presented on the left side of the screen, and Cookie Monster was always presented on the right side of the screen. The correct answer (larger array) was presented equally on the left and right sides such that the number of correct answers was equal across the characters. To control for non-numerical factors, on half of the trials the size of the cookies was matched across arrays (491 pixels^2^) and on the other half, cumulative surface area of the cookies was matched (14726 px^2^) resulting in varied cookie sizes between the arrays, but consistent size within each array. In addition, on half of the trials, the density of the array was larger for the correct answer, and on the other half, the density of the array was smaller for the correct answer.

### 2.6 MR Parameters

Whole-brain BOLD imaging was conducted on a 3-Tesla Siemens MAGNETOM Trio scanner with a 12-channel head coil at the Rochester Center for Brain Imaging. High-resolution structural T1 contrast images were acquired using a magnetization prepared rapid gradient echo (MP-RAGE) pulse sequence at the start of each session [TR = 2530 ms, TE = 3.44 ms flip angle = 7 degrees, FOV = 256 mm, matrix = 256 × 256, 192 or 176 (depending on head size) 1 × 1 × 1 mm sagittal left-to-right slices].

An echo-planar imaging pulse sequence was used for T2* contrast (TR = 2000 ms, TE = 30 ms, flip angle = 90 degrees, FOV = 256 mm, matrix 64 × 64, 30 axial oblique slices, parallel to the AC-PC plane, voxel size = 4 × 4 × 4 mm). For children, an additional sequence was simultaneously acquired with online, retrospective motion correction. Inclusion of data from children was determined by movement measured from the raw, uncorrected sequences, and analyses were conducted on the sequences acquired with online motion correction.

### 2.7 Cognitive Assessments

After the primary imaging session, children completed a behavioral session to evaluate number and letter skills. Number skills were evaluated in three ways to assess 1) rote recitation of the count list, 2) knowledge of the cardinality principle, and 3) the ability to work with and manipulate symbolic and nonsymbolic numbers. To measure memorized counting, children were asked to recite the count list as high as they could up to 100 (“How High?” task based on Huntley-Fenner and Cannon, 2000; Wynn, 1992). Children’s knowledge of the cardinality principle was evaluated using the “Give-N” task up to 10 items (based on Wynn, 1992). Cardinality-principle (CP) knowers were identified as children who could successfully produce sets of 6 items. Subset knowers could successfully produce sets of 1 – 4 items (Le Corre and Carey, 2007; Wynn, 1992). Children’s broader number skills were evaluated using the Test of Early Mathematics Ability Third Edition (TEMA-3, Ginsburg & Baroody, 2003). Finally, we evaluated children’s letter skills using the Kaufman Test of Early Achievement, Letter and Word Recognition Subtest, which tests children’s knowledge of letter names and sounds (K-TEA; Kaufman & Kaufman, 2004). Prior to the Word Processing scan, children were asked to recite the count list as high as they could (“How High?” task) in order to identify suitable number words for the Known and Unknown number word blocks.

### 2.8 Preprocessing

fMRI data were analyzed using BrainVoyager (Goebel et al., 2006) and in-house scripts drawing on the BVQX toolbox (http://support.brainvoyager.com/available-tools/52). The first 2 volumes of functional data in each run were discarded prior to analysis. Preprocessing consisted of slice scan time correction (cubic spline interpolation), motion correction with respect to the first volume in the first run, and linear trend removal in the temporal domain (cutoff: 2 cycles within the run). Functional data were registered to high-resolution anatomy on a participant-by-participant basis in native space. Echo-planar and anatomical volumes were then transformed into Talairach space (Talairach & Tournoux, 1988). Data were normalized into Talairach space by first aligning images with the stereotactic axes and then transforming them to the Talairach grid using a piecewise affine transformation based on manual identification of anatomical landmarks. Analyses were performed on preprocessed data in Talairach space. A Gaussian spatial filter was applied to each volume of functional data at 1.5 voxels (6 mm) FWHM. Average framewise displacement (FD, (Grill-Spector et al., 2008; Power et al., 2012)) was regressed for each child participant using their volume-by-volume realignment parameters. This controls for any increases in signal intensity due to volume-to-volume changes in motion.

### 2.9 Analyses of fMRI Data: General Linear Models

Random effect analyses were used to analyze the group data. Functional data were analyzed using the general linear model. Experimental events were convolved with a standard dual gamma hemodynamic response function. All models included 6 regressors of no interest that corresponded to the motion parameters obtained during preprocessing, and the word processing and numerosity discrimination data included an additional regressor of no interest that corresponded to the button presses. The word processing data were modeled with 5 regressors of interest corresponding to miniblocks of Known and Unknown number and color words and the miniblocks containing catch trials. The numerosity discrimination data were first modeled with one regressor corresponding to the blocks of numerosity discrimination trials and were then modeled with two regressors of interest corresponding to the easy (0.25-ratio) and difficult (0.6-ratio) mini-blocks.

The natural counting data were modeled in two ways. First, we modeled the data with 4 regressors of interest, including the main effect of counting sequences and the main effect of alphabet sequences. In addition, a parametric regressor was included in the model for each condition. There was one parametric regressor across all number word stimuli in the run, and another parametric regressor across all alphabet stimuli in the run. The parametric regressor for counting modeled a linear increase in activation with the magnitude of the number word, that is, each number word was given a weight equivalent to its position in the count list, which also corresponds its magnitude (e.g., “five” was always given a weight of 5, “twenty” was always given a weight of 20, and “seventy-five” was always given a weight of 75, regardless of where it fell in the sequences within each block). The parametric regressor for the alphabet modeled a linear increase in activation with the position of the letter in the sequence (e.g., “C” was always given a weight of 3 and “Y” was always given a weight of 25). Second, we modeled the data with 3 regressors of interest corresponding to number words within children’s counting ranges (“Known” numbers), number words outside of children’s counting ranges (“Unknown” numbers), and alphabet sequences. Children’s counting ranges were determined by asking the children to count as high as they could. For this analysis, we only considered children who could not count to 100 to ensure that each individual had data in both conditions (n = 34 children).

### 2.10 Analyses of fMRI Data: Intersubject Correlations

In a third analysis of the Natural Counting data, intersubject correlations (Hasson et al., 2004) were calculated between children and adults using inhouse scripts in MATLAB 2014B. Intersubject correlations were performed by correlating the full preprocessed timecourses (all 567 volumes) in corresponding voxels between children’s and adults’ whole brain images. Each child’s functional data was correlated with that of each adult (and averaged) to produce an r-map for each child representing the timecourse similarity at each voxel to adults. This image, which can be thought of as how “adult-like” each child’s functional timecourse appears, is referred to as a map of “neural maturity” for this task (Cantlon & Li, 2013). Whole-brain correlations between children’s neural maturity and their number skills (TEMA-3) were calculated using custom scripts that draw on BVQX toolbox in MATLAB. These intersubject correlation maps were previously analyzed as part of a larger study on gender differences and similarities in numerical processing in early childhood (Kersey et al., 2019).

### 2.11 Analyses of fMRI Data: Thresholding

For all whole-brain analyses, results were considered significant at voxel-wise p < 0.01. Whole-brain maps were also cluster corrected to a family-wise p < 0.05 using the BrainVoyager statistical cluster threshold estimator plugin in order to account for multiple comparisons (unless noted for visualizations). Minimum t-values and r-values for each analysis are listed with the degrees of freedom in the corresponding figures. Cluster size, peak coordinates, and peak t-values and r-values are reported for significant regions in tables (Supplement 3) corresponding to each set of analyses in the main text.

### 2.12 Analyses of fMRI Data: Region of Interest Analyses

Region of interest (ROI) analyses were conducted using BrainVoyager, MATLAB 2014B, and R via RStudio. ROIs were defined using independent data from hypothesis tests. The numerosity ROIs used in Figure 3 and Supplement 7 were created in BrainVoyager by drawing a 6mm sphere around the peak voxel in children’s left and right IPS, left and right inferior frontal gyrus (IFG), and anterior cingulate cortex (ACC) as defined by the contrast of (numerosity task+). Single-subject beta values were extracted for the easy and difficult ratio trials from the numerosity task using the BVQX toolbox in MATLAB 2014B. Correlations were calculated using the “cor.test” function in R.

**Figure 1.**
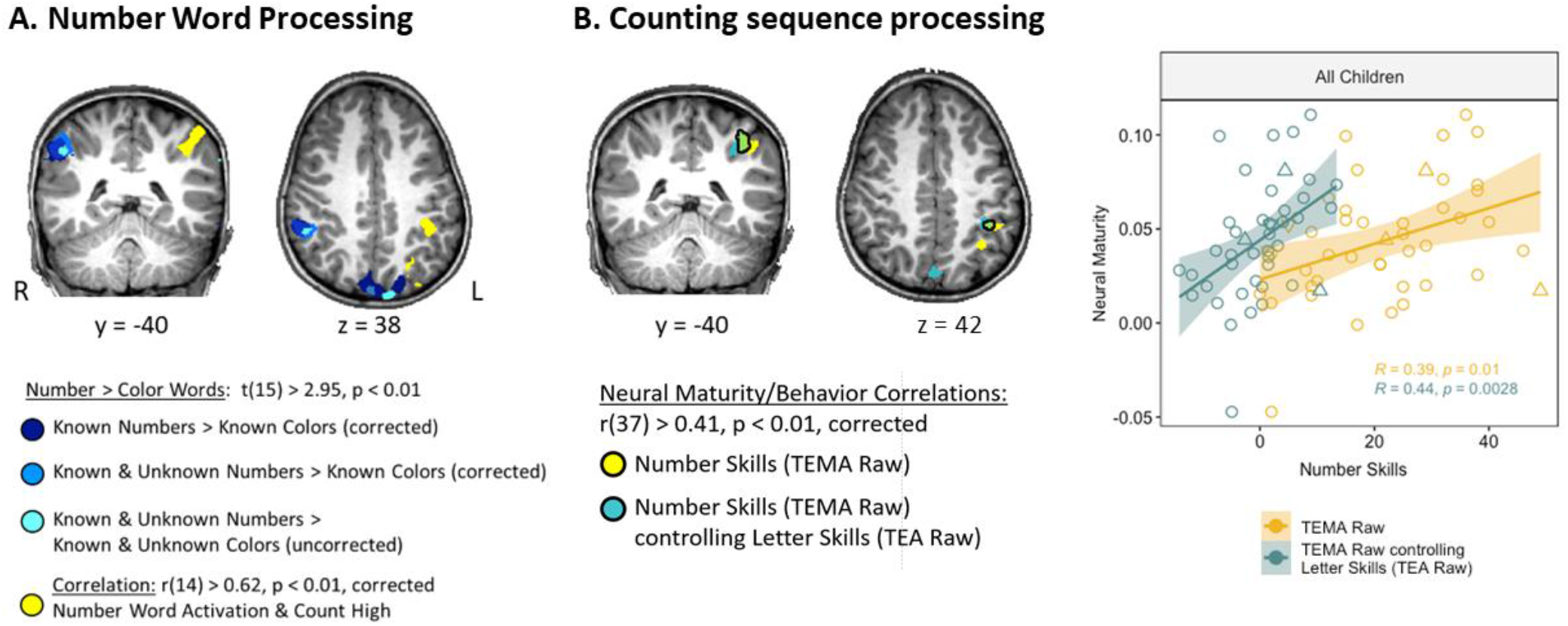
(A) Recruitment of regions for number word processing. (B) Recruitment of regions for counting sequence processing during the Natural Counting paradigm. Left: Regions of left IPS where there was a significant correlation between children’s neural maturity (similarity to adults) and behavioral number skills. Right: Correlation between children’s neural maturity and behavioral number skills, with (green) and without (orange) controlling for behavioral letter skills (TEA Raw).

**Figure 2.**
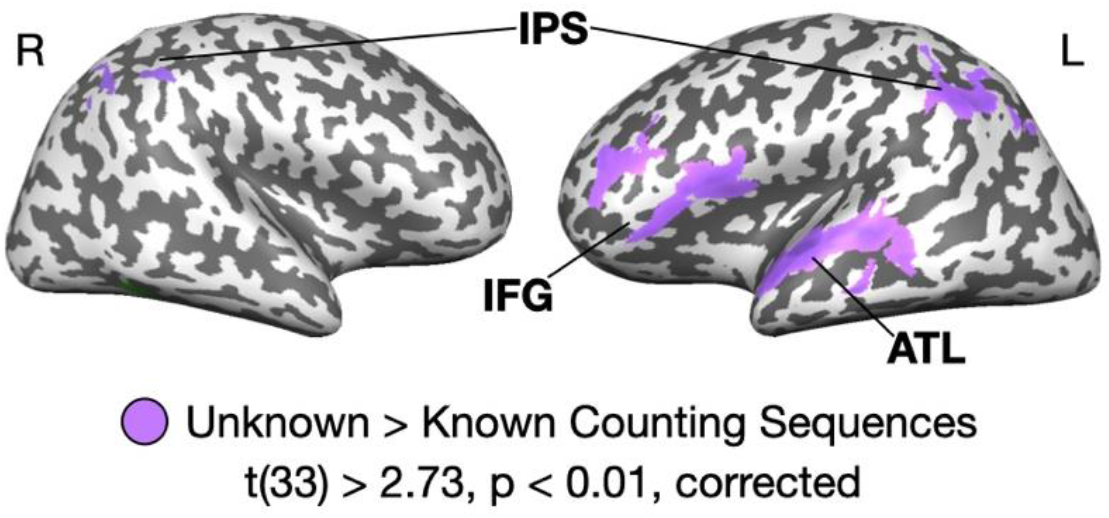
Neural representations while listening to unknown counting sequences compared to known counting sequences. Results are shown at voxel-wise p < 0.01, corrected with an applied cluster threshold of 25 mm^3^. Peaks reported in Talairach Coordinates: IPS (intraparietal sulcus): Left IPS x = -41, y = -58, z = 37, Right IPS x = 43, y = -51, z = 51; IFG (inferior frontal gyrus): x = -45, y = 7, z = 9; ATL (Anterior Temporal Lobe) Peaks in Superior Temporal Gyrus (STG) x = - 56, y = -8, z = 1 and Middle Temporal Gyrus (MTG): x = -61, y = -24, z = -2). A full list of regions is presented in Supplement 3, Table S5.

**Figure 3.**
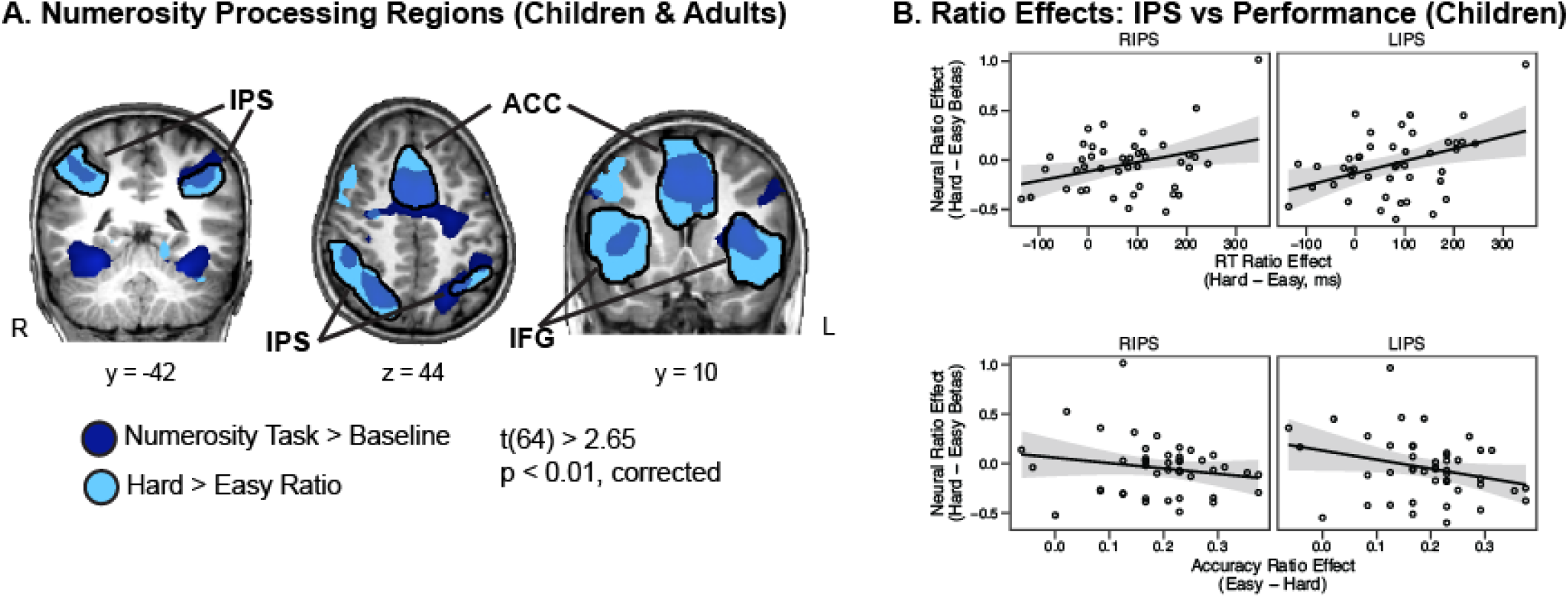
Numerosity Processing. (A) Regions of the brain recruited during the numerosity task (dark blue) and those that showed a ratio effect (light blue). A full list of coordinates and regions is presented in Supplement 3, Table S6. (B) Correlations between children’s neural ratio effects in the IPS and their performance on the numerosity comparison task. Results for the IFG and the ACC are shown in Supplement 7, Figure S4.

For the Bayes Factor ROI analyses presented in Figure 6 and Supplement 9, ROIs consisted of the full numerosity processing ROIs as identified by a BF10 ≥ 3 for the neural ratio effect (Hard > Easy trials) in adults only. BF10 values were then extracted from those entire regions for 3 other neural contrasts (child numerosity processing (children task+), parametric effect of counting, and unknown > known counting sequences) using the BVQX toolbox in MATLAB 2014B.

**Figure 4.**
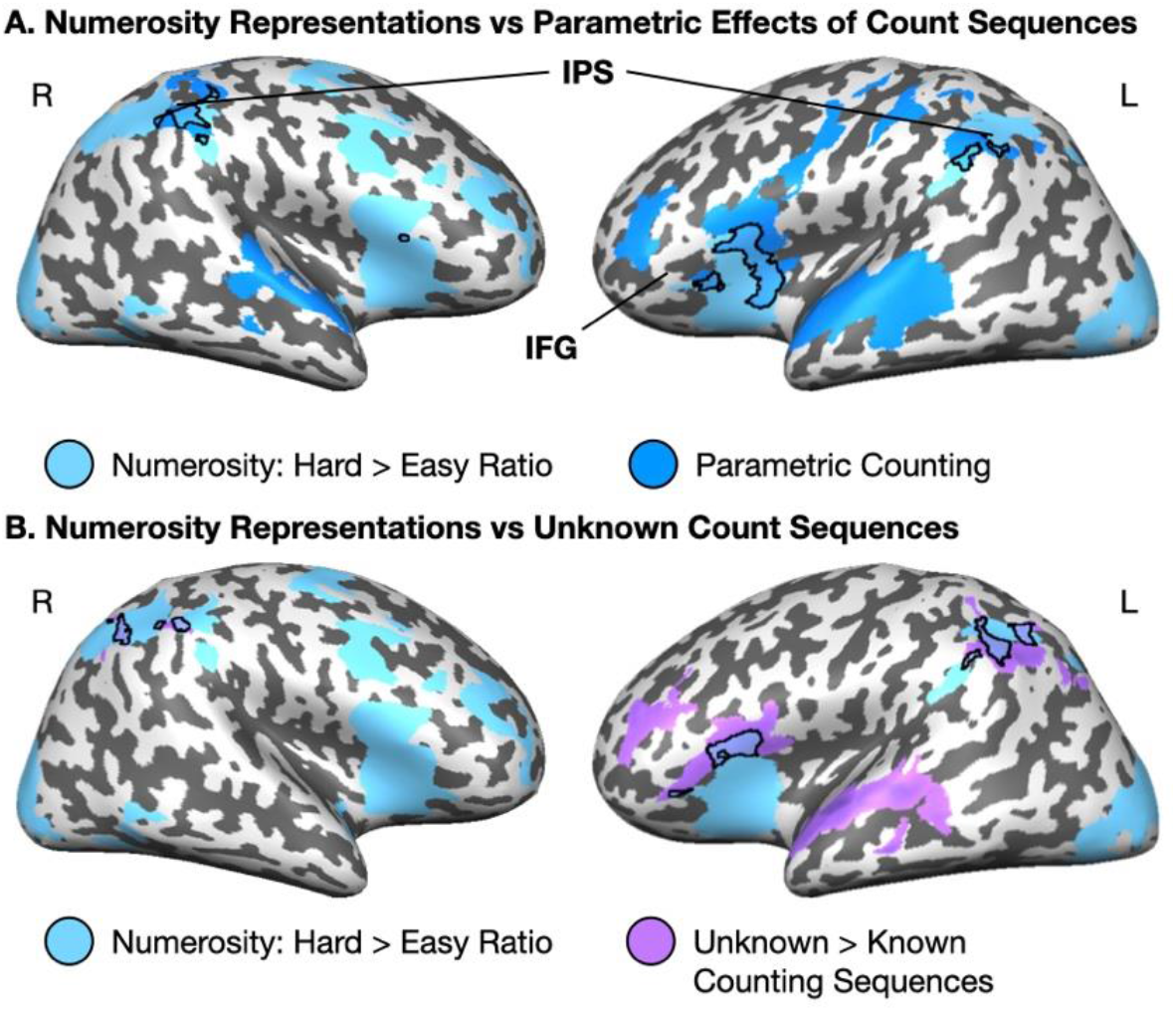
Functional overlap between numerosity representations (light blue) and (A) parametric counting representations (dark blue) and (B) representations of unknown count sequences (purple). Overlap between numerosity and counting in each panel is outlined. All three effects overlapped in left IPS (TAL: -41, -53, 43), right IPS (TAL: 42, -46, 50), and left IFG (TAL: -46, 8, 8).

**Figure 5.**
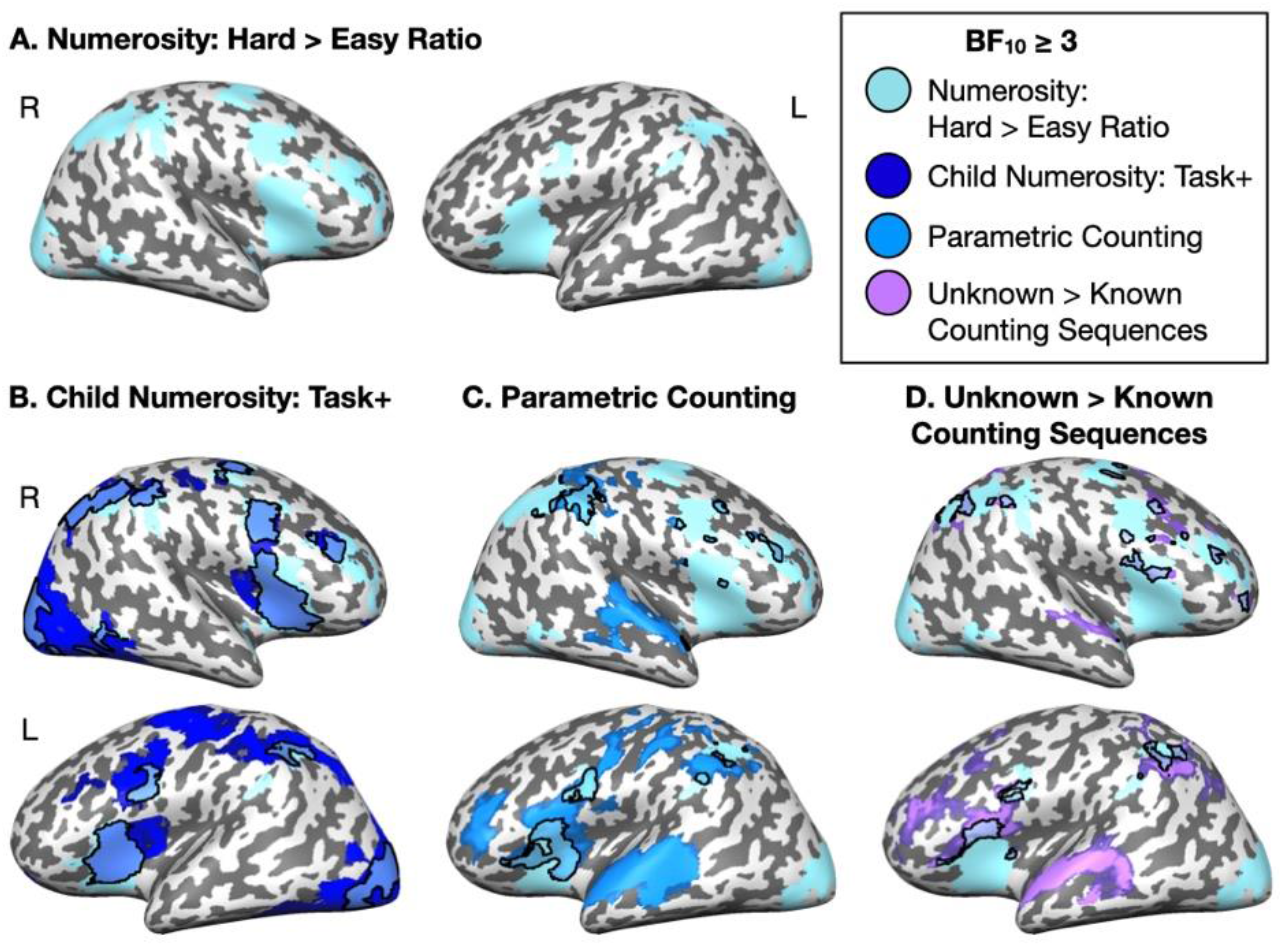
Bayes Factors > 3 for key contrasts: (A) Mature Numerosity Processing: Hard Ratio > Easy Ratio in children and adults, (B) Child Numerosity Processing: Hard Ratio + Easy Ratio in children, (C) Parametric Effect of Counting, (D) Unknown vs Known Counting Sequences. The overlap between Mature Numerosity Processing and the main effects in panels B-D are outlined. All maps are displayed with an arbitrary applied cluster threshold of 25 mm^3^.

**Figure 6.**
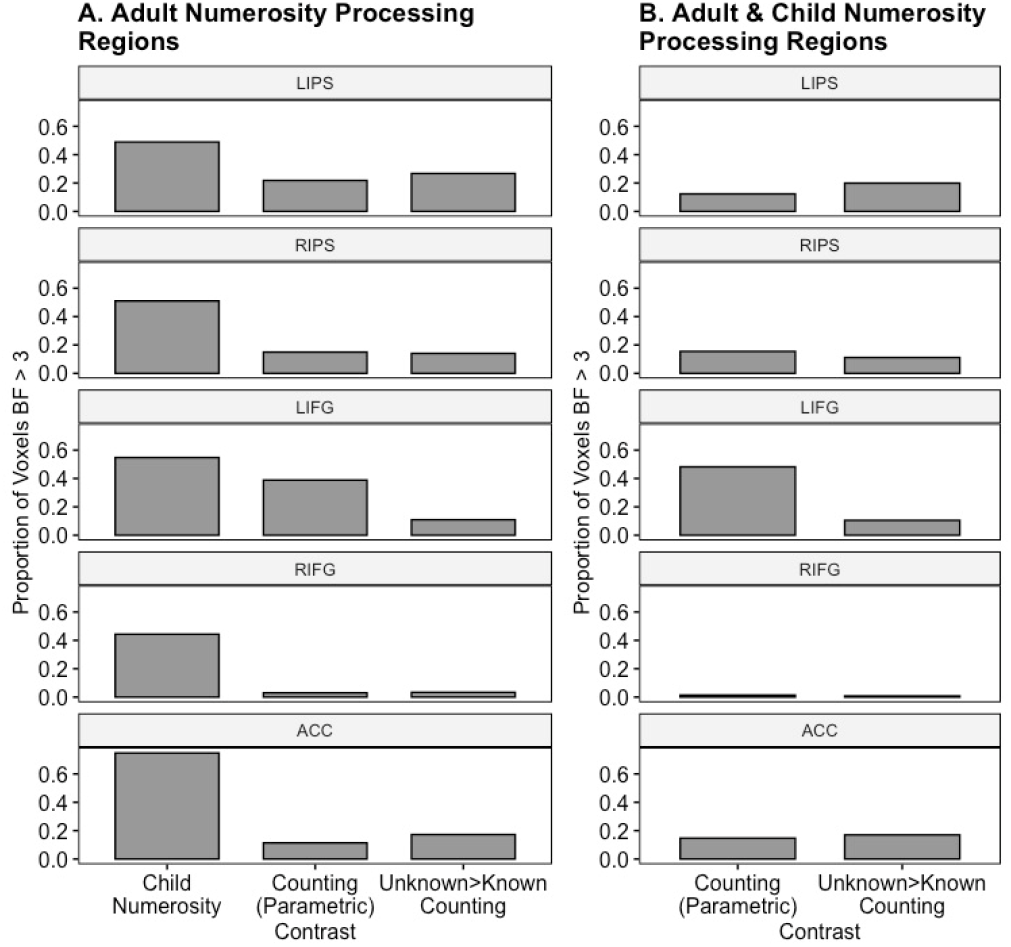
Strength of effects as measured by Bayes Factors within regions that show substantial effects of the mature ratio processing effect (BF10 ≥ 3 for hard > easy ratio across all participants). (A) Proportion of adult numerosity processing voxels with BF10 ≥ 3 in the left and the right IPS, left and right IFG, and ACC for the child numerosity effect, parametric counting effect, and Unknown > Known counting effect. (B) Proportion of adult and child numerosity processing voxels with BF10 ≥ 3 for the parametric counting effect and Unknown > Known counting effect.

## 3. Results

### 3.1 Neural representations of spoken number words in early childhood

First, we identified regions of the developing brain that showed a preference for number words over color words during the Word Processing task. Three GLM analyses were conducted to compare 1) known number words to known color words, 2) all number words to known color words, and 3) all number words to all color words. The first comparison of known number words to known color words represents the strictest identification of number word representations because it ensures that children can distinguish between words that signify numbers vs those that signify colors. In line with our a priori hypotheses, regions of the right IPS showed greater activation during known number word processing than during known color word processing (Figure 1A, dark blue, Supplement 3, Table S2; Known Number Words > Known Color Words, t(15) > 2.95, voxel-wise p < 0.01, cluster corrected to p < 0.05, cluster threshold = 52 voxels). The second comparison tested whether regions that represented number words that children could count to also represented number words that children have not yet learned but might recognize as number words (Sarnecka & Gelman, 2004; Wang & Feigenson, 2019). For example, a child who knows that “two” and “seven” are number words might recognize that “seventy-two” is also a number word. The right IPS represented both known and unknown number words over and above known colors, suggesting that this region processes number words, whether known or unknown, differently than non-number words (Figure 1A, green, Supplement 3, Table S2; Known Number Words + Unknown Number Words > Known Color Words (balanced), t(15) > 2.95, voxel-wise p < 0.01, cluster corrected to p < 0.05, cluster threshold = 51 voxels). An alternative possibility is that the same regions that represent number words also represent any words that children have not acquired. The last comparison tested whether a preference for number words persisted when unknown color words were considered in the analysis. The right IPS continued to show a preference for number word processing when unknown color words were added to the model, though the number-preferring effect was slightly weaker (Figure 1A, light blue, Supplement 3, Table S2; Known Number Words + Unknown Number Words > Known Color Words + Unknown Color Words, t(15) > 2.95, voxel-wise p < 0.01, uncorrected). Nonetheless, the right IPS showed a preference for number words across all comparisons indicating a degree of specialization for processing number words even before children can recite the entire count list.

To determine if any regions showed patterns of change related to number word acquisition, we tested for correlations between children’s abilities to recite the count list and their number word preferring activation (as defined by the neural response to the Known & Unknown Numbers > Known & Unknown Colors contrast). This effect was found in the left IPS (Figure 1A, yellow, Supplement 3, Table S2; r(14) > 0.62, voxel-wise p < 0.01, cluster-corrected to p < 0.05, cluster threshold = 48 voxels) suggesting that as children get better at counting they show more specificity for number word processing over color word processing in the left IPS.

Taken together, we find that in 3- to 5-year-olds the right IPS shows a preference for representing number words over color words, and the left IPS shows this effect as children’s knowledge of the count list improves. These findings are consistent with the prediction that numerosity representations in the IPS interface with count words during the acquisition of verbal counting in children.

### 3.2 Counting ability and neural development

Previous work suggests that parietal cortex represents patterns, sequences, and other ordinal information (Fias et al., 2007; Franklin & Jonides, 2009; Marshuetz et al., 2006; Matejko et al., 2018; Pariyadath et al., 2012; Turconi & Seron, 2002; Wang et al., 2015). To investigate the possibility that the IPS represents ordinality, we compared neural representations of the count list to neural representations of the alphabet, which is another ordinal list that children begin to memorize around the same time as the count list. If the localization of number word processing to parietal cortex is related to the ordinal information contained in number words, then counting and alphabetic sequences should recruit the same neural substrates within parietal cortex. Alternatively, if number word processing in parietal cortex is sensitive to the semantic property of quantity, then number sequences should dissociate from the alphabet in parietal cortex.

Regions that represented sequences in early childhood were identified by testing for a parametric modulation of activation with the ordinal position of the number word or letter during the spoken sequences, which would indicate a sensitivity to the sequential aspects of the stimuli. We found that ordinal representations of both counting and alphabet sequences were evident in the left and right inferior frontal gyri and the left and right anterior temporal lobe (Supplement 3, Table S3; Supplement 5, Figure S2; t(42) ≥ 2.70, voxel-wise p < 0.01, cluster-corrected to p < 0.05, cluster thresholds = 102 voxels for counting sequences, 75 voxels for alphabet sequences). In contrast, neural activation in the IPS showed a different pattern. Although overall levels of activation were similar across counting and alphabet sequences (mean-normalized percent signal change LIPS: counting = -0.009, alphabet = 0.006, t(42) = 0.78, p = 0.43; RIPS: counting = -0.003, alphabet = 0.002, t(42) = 0.48, p = 0.63; see Supplement 3, Table S3 and Supplement 5, Figure S2B for whole-brain results), the parametric effect of sequence processing differed across stimuli types. Counting sequences, but not alphabet sequence, were represented in the left and the right IPS, and alphabet sequences were uniquely represented in the right prefrontal cortex (middle and superior frontal gyri) and the right inferior parietal lobule. This suggests that the neural representations of counting in the IPS are specifically sensitive to semantic properties of number words.

To determine whether the IPS is important for the acquisition of counting, we first tested for a relation between children’s number skills (TEMA-3) and the “neural maturity” of their sequence processing during the Natural Counting paradigm. Neural maturity represents the degree to which the children showed “adult-like” temporal patterns of neural activity during the natural counting task and was assessed by conducting intersubject timeseries correlations between neural activity from children and adults during the natural counting task. Regions of the left IPS and bilateral posterior parietal cortex showed greater neural maturity during numerical sequence processing in children who had more advanced number skills (r(37) > 0.41, voxel-wise p < 0.01, cluster-corrected to p < 0.05, cluster threshold = 21 voxels; see Figure 1B and Supplement 3, Table S4). This effect persisted when controlling for letter skills (r(37) > 0.41, voxel-wise p < 0.01, cluster-corrected to p < 0.05, cluster threshold = 22 voxels; see Figure 1B and Supplement 3, Table S4), indicating that the neural maturity of this region for sequence processing is related to numerical development.

Next, we tested for developmental change related to counting acquisition by directly comparing sequences within a child’s counting range (“Known” counting sequences) and sequences outside a child’s counting range (“Unknown” counting sequences). Regions that are involved in the acquisition of number words will likely respond to Unknown number words, whereas regions that represent number words after acquisition will show greater activation for Known number words. A whole-brain contrast revealed that the IPS was more strongly recruited for Unknown counting sequences, suggesting a role for the IPS in counting acquisition (Figure 2; see Supplement 3, Table S5 for a list of significant regions). Unknown counting sequences also more strongly recruited regions of the left IFG and superior and lateral left ATL spanning the superior and middle temporal gyri (S/MTG; Figure 2). To ensure that this effect was not driven by linguistic complexity of larger number words, we tested for a parametric effect of the number of syllables in the stimuli presented in both the word processing and natural counting tasks. Syllable effects were evident in the bilateral superior temporal gyrus (each map entered at voxel-wise p < 0.01, uncorrected; word processing: t(15) > 2.95, natural counting: t(42) > 2.70). Importantly, there was no syllable-related activation in the IPS (Supplement 6, Figure S3) suggesting that the Unknown count word effect does not reflect increased complexity of larger number words. Together, these results show that the IPS not only represents number words, but is also recruited for processing Unknown number words, suggesting a role in the acquisition of number word meanings.

### 3.3 Neural processing of numerosity

If number words acquire meaning via primitive perceptual mechanisms for numerosity processing in parietal cortex, there should be functional overlap between numerosity representations and representations of counting in children who are learning to count, and this overlap should be evident for number words that children are still learning (i.e., Unknown number words). To test this prediction, we first identified neural representations of numerosity by conducting a t-test of Numerosity Task > Baseline across children and adults. Consistent with previous studies (e.g., Cantlon & Li, 2013; Emerson & Cantlon, 2012), we identified a set of regions in bilateral IPS, bilateral IFG, ACC, and bilateral occipital cortex that children and adults recruited during the numerosity discrimination task (Figure 3A, dark blue; Supplement 3, Table S6).

Then, we tested for a main effect of comparison ratio in a 2 (age-group: children, adults) x 2 (ratio: difficult – 0.6 ratio, easy – 0.25 ratio) ANOVA on neural amplitudes from the numerosity discrimination task. A main effect of ratio (stronger activation for difficult trials) was revealed in bilateral IPS, bilateral IFG, ACC, and bilateral occipital cortex (Supplement 3, Table S6; t(67) > 2.65, voxel-wise p < 0.01, cluster-corrected to p < 0.05, cluster threshold = 91 voxels). Consistent with previous literature (Ansari & Dhital, 2006; Lussier & Cantlon, 2017), the ratio effect was stronger in adults than in children. In children, the neural ratio effect was related to task performance. The neural ratio effect in the IPS was positively correlated with children’s response time ratio effect (RT ratio effect: RT for difficult trials – RT for easy trials; neural ratio effect: beta for difficult ratios – beta for easy ratios; Figure 3B; right IPS: r(42) = 0.34, t(42) = 2.35, p = 0.024; left IPS: r(42) = 0.40, t(42) = 2.63, p = 0.012). Children’s accuracy showed a similar pattern but a marginal effect (Accuracy ratio effect: accuracy for easy trials – accuracy for difficult trials; Figure 4B; right IPS: r(43) = -0.19, t(43) = -1.26, p = 0.215; left IPS: r(43) = -0.28, t(43) = -1.95, p = 0.058). Similar patterns were identified in the frontal regions of the numerosity network (Supplement 7). In sum, children and adults engage canonical number processing regions for numerosity comparisons. In children, the strength of the neural ratio effect was modulated by task performance, suggesting that, like behavioral precision, neural precision for numerosity representations is still developing.

### 3.4 Functional Overlap between Neural Representations of Counting and Numerosity Processing

Finally, we compared neural representations of counting to neural representations of numerosity processing. Because children’s performance on the numerosity comparison task influenced their neural ratio effects, here we used the main effect of ratio as an index for numerosity processing. Consistent with our predictions, we found that neural representations of counting and numerosity overlapped in the IPS (Figure 4). Critically, this overlap was also evident for number words that children had not learned to count to (Figure 4B), and a similar pattern was evident in the subset of children who had not yet acquired the cardinality principle (Supplement 8). A region in the left IFG also showed overlap among neural representations of numerosity, counting sequences, and Unknown counting sequences (Figure 4).

To quantify the strength of evidence for the neural overlap between numerosities and number words in children, we conducted whole-brain Bayes Factor analyses for four neural contrasts: parametric effect of counting, Unknown > Known number words, numerosity processing in children (hard ratio + easy ratio), and mature numerosity processing (hard ratio > easy ratio in children and adults). Complementing the frequentist results, we found substantial evidence of early childhood numerosity and counting representations in the IPS (BF10 ≥ 3). The mature numerosity effect showed substantial and strong evidence within the regions identified by the frequentist analysis, including the left and right IPS and the left and right IFG (Figure 5A). Similar regions showed strong evidence for numerosity processing in young children alone (Figure 5B). As in the frequentist analyses, there was substantial evidence of an increase in activation with ordinal position in the count list within the right and the left IPS, the right and the left ATL, and the left IFG (Figure 5C). Similarly, Unknown count words showed substantial evidence of stronger activation than Known count words within the left and the right IPS, the left ATL, and the left IFG (Figure 5D). Smaller clusters within the right ATL and the right IFG also showed substantial evidence of this effect using a Bayes Factor approach. Within the left and the right IPS, voxels showed substantial evidence of both numerosity processing and the counting effects.

To further quantify these effects, we conducted a region-of-interest analysis within the primary regions of the numerosity processing network. Regions within the left and right IPS, the left and right IFG, and the ACC that showed a substantial effect of numerosity processing in adults (BF10 ≥ 3; hard > easy ratio) were defined as regions of interest. Data from adults were used to identify regions of interest to maintain independence from the data from children. We extracted Bayes Factors within these regions for the three other contrasts (Child Numerosity Processing, Parametric Effect of Counting, Unknown > Known Counting) and evaluated the proportion of voxels within the adult numerosity regions that showed a substantial effect for each contrast (Figure 6A). That number represents the proportion of numerosity processing voxels that show functional overlap with early childhood representations of number.

We found that 50% of IPS voxels showed substantial evidence of numerosity processing in children (LIPS: 49%, RIPS: 51%, LIFG: 55%, RIFG: 44%, ACC: 75%), suggesting strong consistency between the child and adult numerosity effects. Fewer voxels showed overlap with the counting effects. 15% - 22% of IPS voxels that showed an effect of numerosity processing also showed a substantial effect of parametric counting in children (LIPS: 22%, RIPS: 15%, LIFG: 39%, RIFG: 3%, ACC: 11%), and 14% - 27% of IPS voxels showed a substantial effect of Unknown vs Known count sequences in children (LIPS: 27%, RIPS:14%, LIFG: 11%, RIFG: 3%, ACC: 17%). Of the IPS voxels that showed functional overlap, half showed strong effects of parametric counting (BF10 ≥ 10, LIPS: 61%, RIPS: 54%, LIFG: 48%, RIFG: 71%, ACC: 36%), and approximately one third of IPS voxels showed strong evidence of representations for Unknown count words (see Supplement 9 for more details; Unknown > Known Count Sequences: BF10 ≥ 10, LIPS: 37%, RIPS: 31%, LIFG: 49%, RIFG: 19%, ACC: 51%).

Finally, we examined the strength of evidence for counting effects within the voxels that showed substantial effects of both adult numerosity processing and child numerosity processing (BF10 ≥ 3 for both numerosity effects; Figure 6B). Within these strictly defined numerosity processing voxels, 12%-15% of IPS voxels showed an effect of parametric counting (LIPS: 12%, RIPS: 15%, LIFG: 48%, RIFG: 1%, ACC: 15%), and 11% - 20% of IPS voxels showed an effect of Unknown counting sequences (LIPS: 20%, RIPS: 11%, LIFG: 11%, RIFG: 1%, ACC: 17%). This provides evidence of functional overlap in the IPS for 1) numerosity processing in adulthood, 2) numerosity processing in early childhood, and 3) count sequence processing in young counters.

## 4. Discussion

The present experiments tested the prediction that primitive, perceptual representations in parietal cortex help assign meaning to number words. Previous work demonstrated that numerosity representations and symbolic number word representations are linked by 8 years of age (Lussier & Cantlon, 2017). However, the role of numerosity representations in the acquisition and development of symbolic number representations during childhood is debated (Carey & Barner, 2019; Wilkey & Ansari, 2020). Some studies suggest that numerosity representations do not play a role in number word acquisition because children cannot successfully map from quantities to number words until after they have acquired the first four or five number words (Le Corre & Carey, 2007; Lipton & Spelke, 2005). Other studies show that children can map in the opposite direction (i.e. from number words to quantities) even earlier than this (Gunderson et al., 2015; Odic et al., 2015; Opfer et al., 2010; Wagner & Johnson, 2011). In the current set of experiments, we showed that early perceptual numerosity mechanisms interface with language mechanisms as children process spoken number words and count sequences, even words for which they do not yet know the precise meanings. These findings suggest that non-symbolic numerosity representations play an important role in the acquisition of symbolic number representations.

Previous research with older children and adults implicates the IPS, the IFG, and the fronto-temporal language network as candidates for number word acquisition (Ansari et al., 2005; Cantlon et al., 2009a; Diester & Nieder, 2007; Lussier & Cantlon, 2016; Lyons & Ansari, 2009; Nieder, 2009; Rivera et al., 2005), but previous studies were unable to test the acquisition of new number words and concepts because they only tested older participants who already knew all the words and concepts. Here, 3- to 5-year-old children’s representations of count words evoked activation in the IPS, the IFG, and the ATL. The IPS region is important because it overlapped in part with children’s activation to numerosities. Although the overlap was a small portion of the total activation in response to numerosities and to number words, the Bayesian analyses suggest that when this overlap is present, the effects are quite reliable. Furthermore, there was evidence of overlap despite differences in the tasks and demands. In the natural counting task, the numeric information was presented auditorily, and the task was passive listening. In contrast, in the numerosity discrimination task, numeric information was presented visually, and the task was active perceptual discrimination. That two very different tasks engaged overlapping neural substrates in the IPS lends support to the hypothesis that the IPS plays a developmental role in number word semantics.

Across two number word tasks, behavioral acquisition of number words was systematically related to neural activity in the left IPS. Children who could count higher showed a greater preference there for number words over color words, and children with stronger numerical concepts showed more adult-like patterns of count sequence processing, even after controlling for letter skills. Importantly, this cannot be attributed to a lack of response to unfamiliar number words because the left and right IPS were recruited more strongly for counting sequences beyond children’s count ranges compared to counting sequences within children’s count ranges. Even children who had not acquired the cardinality principles showed strong recruitment of the left IPS for number words that they had not learned to count. However, this effect was only identified when number words were presented in counting sequences. When number words were presented randomly as in the ‘Word Processing’ task, no regions showed differences between Known and Unknown number words. One possibility is that listening to count sequences allows children to begin to link Unknown number words with their magnitudes better than when those words are presented outside the count list. Together these findings suggest the IPS supports early representations of number words with the right IPS playing a strong role throughout number word acquisition while the left IPS matures gradually to support the meanings of number words. This interpretation is consistent with previous work that shows early maturation of the right IPS for numerical processing and developmental change in the left IPS related to symbolic mathematics development (Ansari et al., 2006; Bugden et al., 2012; Cantlon et al., 2006; Cantlon & Li, 2013; Emerson & Cantlon, 2014; Holloway & Ansari, 2010; Hyde et al., 2010; Kersey & Cantlon, 2017b; Rosenberg-Lee et al., 2011; Vogel et al., 2015). One hypothesis is that the left hemisphere is more strongly related to math skills because closely related skills, such as reading and speech, are also left lateralized. Overall the findings in the IPS support theories that symbolic mathematics is grounded in primitive, numerosity representations (e.g., Dehaene & Cohen, 2007) and suggest that the functional overlap seen in older children and adults reflects the origin rather than the outcome of long-term mathematical training (Cantlon et al., 2009a; Holloway & Ansari, 2010; Lussier & Cantlon, 2017; Piazza et al., 2007).

The IFG and the left ATL also represented count sequences. Like the IPS, the IFG was recruited for both numerosity processing and for representing count sequences, and the left IFG showed stronger recruitment for unknown number words compared to known ones. Unlike the IPS, the IFG also represented alphabet sequences. Representations of alphabet sequences suggest that this region may function in a domain-general manner by representing ordinal information of sequences (Wang et al., 2015). The left ATL did not distinguish counting and alphabet sequences but showed greater recruitment for unknown counting sequences compared to known ones. The left ATL also showed strong activation to spoken number and color words, suggesting a linguistic function in the acquisition of number words. This conclusion is consistent with previous research showing that the left ATL functions as part of a lexical-semantic network during the acquisition of word meanings (Kotz et al., 2002).

The current study shows that children’s numerosity representations in the IPS along with the lexical-semantic language network (i.e., IFG and ATL) are involved in representing number words during the acquisition of counting. Functional overlap between perceptual numerosity representations and symbolic number word representations in the IPS in the youngest counters (3- to 5-year-olds) is consistent with theoretical accounts of cultural recycling (e.g., Dehaene & Cohen, 2007) and symbol grounding (e.g., Barsalou, 1999) in the developing brain, which predict that the acquisition of formal mathematics builds on evolutionarily-primitive, perceptual numerosity representation in the IPS. These data provide a novel empirical demonstration of integration between evolutionarily recent language systems and evolutionarily primitive perceptual systems in the acquisition of children’s counting.

## Acknowledgements

The authors would like to thank Courtney Lussier, Kelsey Csumitta, Jill Schwartz, Adina Levitt, Rebecca Lawrence, and Pat Weber for their help with data collection.

## Ethics Statement

All research was approved by the University of Rochester Research Subjects Review Board. Written informed consent was obtained from adult participants and parents of the child participants.

## Descriptive Titles of Electronic Supplementary Material

Supplement 1: Color word stimuli used in the ‘Word Processing’ task

Supplement 2: Movie paradigm shown to participants during the ‘Natural Counting’ task

Supplement 3: Tables of results

Supplement 4: Additional results from the ‘Word Processing’ task

Supplement 5: Children’s neural responses during the ‘Natural Counting’ task

Supplement 6: Neural processing of word complexity

Supplement 7: Correlations between neural ratio effect and behavioral ratio effects in children

Supplement 8: Unknown count word processing in subset knowers

Supplement 9: Distribution of BF10 values in voxels that represent numerosity and counting

## Footnotes

1 The first child and the first adult participants who participated in this study completed this task as a Go/No-Go task and were instructed to press the button whenever Cookie Monster had the most cookies.

