## Supplemental Information for "Emergence of counting in the brains of 3- to 5-year-old children"

**Supplement 1**

**Table S1.** Color word stimuli and their numbers of syllables presented in the word listening fMRI paradigm.

| Known Color Words | | | |
|---|---|---|---|
| Word | Syllables | Word | Syllables |
| Black | 1 | Blackish-Blue | 3 |
| Blue | 1 | Blueish-Green | 3 |
| Brown | 1 | Dark Orange | 3 |
| Gold | 1 | Dark Purple | 3 |
| Gray | 1 | Dark Yellow | 3 |
| Green | 1 | Grayish-Brown | 3 |
| Peach | 1 | Greenish-Blue | 3 |
| Pink | 1 | Greenish-Brown | 3 |
| Red | 1 | Light Purple | 3 |
| White | 1 | Light Yellow | 3 |
| Dark Brown | 2 | Orangish-Red | 3 |
| Dark Red | 2 | Reddish-Pink | 3 |
| Light Blue | 2 | Purpleish-Blue | 4 |
| Light Green | 2 | Greenish-Yellow | 4 |
| Light Pink | 2 | Orangish-Yellow | 4 |
| Orange | 2 | Pinkish-Purple | 4 |
| Purple | 2 | Reddish-Purple | 4 |
| Silver | 2 | Yellowish-Brown | 4 |
| Yellow | 2 | Yellowish-Orange | 5 |

| Unknown Color Words | | | |
|---|---|---|---|
| Word | Syllables | Word | Syllables |
| Mauve | 1 | Indigo | 3 |
| Bronze | 1 | Lavender | 3 |
| Puce | 1 | Magenta | 3 |
| Teal | 1 | Navy Blue | 3 |
| Ash Gray | 2 | Ruby Red | 3 |
| Brick Red | 2 | Sea Foam Green | 3 |
| Chartreuse | 2 | Violet | 3 |
| Fuchsia | 2 | Burnt Sienna | 4 |
| Off-White | 2 | Cerulean | 4 |

| | | | |
|---|---|---|---|
| Turquoise | 2 | Mahogany | 4 |
| Amethyst | 3 | Blue Violet | 4 |
| Neon Pink | 3 | Royal Purple | 4 |
| Burgundy | 3 | Mustard Yellow | 4 |
| Burnt Orange | 3 | Wisteria | 4 |
| Ebony | 3 | Cadmium Yellow | 5 |

**Supplement 2**
See additional supplementary video file for a recording of the Movie paradigm shown to participants during the 'Natural Counting' task.

**Supplement 3**

**Table S2.** Word Listening Results.

**All Number & Color Words > Baseline: t(15) > 2.95, p < 0.01, cluster corrected**

| Region | Hemisphere | Size ($mm^3$) | Peak X | Peak Y | Peak Z | t |
|---|---|---|---|---|---|---|
| Anterior Temporal Lobe/Superior Temporal Gyrus | Left | 32366 | -48 | -22 | 1 | 16.67 |
| Anterior Temporal Lobe/Superior Temporal Gyrus | Right | 26736 | 60 | -16 | 1 | 13.70 |
| Inferior Frontal Gyrus/Insula | Left | 7822 | -33 | 11 | 22 | 6.06 |
| Inferior Frontal Gyrus/Insula | Right | 6875 | 42 | 20 | 13 | 5.29 |

**Known Number > Known Color: t(15) > 2.95, p < 0.01, cluster corrected**

| Region | Hemisphere | Size ($mm^3$) | Peak X | Peak Y | Peak Z | t |
|---|---|---|---|---|---|---|
| Intraparietal Sulcus | Right | 1795 | 48 | -40 | 37 | 4.57 |
| Medial Frontal Gyrus | Medial | 7512 | 6 | 62 | 10 | 4.82 |
| Cuneus | Medial | 11336 | 3 | -82 | 37 | 5.35 |
| Occipital Cortex/Cerebellum | Left | 7062 | -27 | -88 | -24 | 4.97 |

**Known & Unknown Number > Known Color: t(15) > 2.95, p < 0.01, cluster corrected**

| Region | Hemisphere | Size ($mm^3$) | Peak X | Peak Y | Peak Z | t |
|---|---|---|---|---|---|---|
| Intraparietal Sulcus | Right | 1759 | 48 | -40 | 37 | 5.05 |
| Medial Frontal Gyrus | Medial | 8094 | 0 | 66 | 25 | 5.23 |

**Known & Unknown Number > Known & Unknown Color: t(15) > 2.95, p < 0.01, uncorrected**

| Region | Hemisphere | Size ($mm^3$) | Peak X | Peak Y | Peak Z | t |
|---|---|---|---|---|---|---|

| | | | | | | |
|---|---|---|---|---|---|---|
| Intraparietal Sulcus | Right | 107 | 45 | -40 | 37 | 3.21 |
| Inferior Parietal Lobule | Left | 108 | -57 | -34 | 28 | 3.34 |
| Postcentral Gyrus | Right | 140 | 12 | -58 | 68 | 3.95 |
| Middle Frontal Gyrus | Right | 2267 | 40 | 59 | 4 | 4.36 |
| Middle Frontal Gyrus | Right | 361 | 30 | 5 | 49 | 3.86 |
| Medial Frontal Gyrus | Medial | 155 | 3 | -7 | 55 | 3.17 |
| Middle Frontal Gyrus | Left | 1303 | -21 | -1 | 46 | 4.71 |
| Inferior Temporal Gyrus | Right | 115 | 54 | -10 | -29 | 4.10 |
| Precuneus | Left | 910 | -9 | -79 | 46 | 4.56 |
| Precuneus | Left | 651 | -3 | -52 | 58 | 3.82 |
| Cuneus | Medial | 791 | 0 | -91 | 29 | 4.38 |
| Occipital Cortex | Right | 209 | 24 | -70 | -2 | 3.64 |
| Occipital Cortex | Right | 124 | 9 | -55 | 1 | 3.62 |
| Middle Occipital Gyrus | Left | 3229 | -30 | -92 | 19 | 5.75 |
| Occipital Cortex | Left | 1468 | -51 | -64 | -17 | 4.23 |
| Occipital Cortex | Left | 230 | -12 | -70 | -2 | 3.61 |
| Occipital Cortex/Cerebellum | Left | 3146 | -9 | -91 | -20 | 4.46 |
| Cerebellum | Right | 159 | 33 | -61 | -26 | 3.56 |
| Caudate | Left | 315 | -18 | -25 | 22 | 3.98 |
| Cingulate Gyrus | Left | 136 | -6 | -22 | 37 | 3.34 |

**Unknown Color > Known Color: $t(15) > 2.95$, $p < 0.01$, cluster corrected**

| Region | Hemisphere | Size ($mm^3$) | Peak X | Peak Y | Peak Z | t |
|---|---|---|---|---|---|---|
| Superior Frontal Gyrus | Left | 1319 | -24 | 53 | 25 | 5.03 |
| Fusiform Gyrus | Left | 1648 | -55 | -4 | -23 | 4.92 |

| Unknown Color > Unknown Number: t(15) > 2.95, p < 0.01, cluster corrected | | | | | | |
|---|---|---|---|---|---|---|
| Region | Hemisphere | Size (mm$^3$) | Peak X | Peak Y | Peak Z | t |
| Middle Frontal Gyrus | Left | 3567 | -18 | 23 | -8 | 5.21 |

| Known Number > Unknown Color: t(15) > 2.95, p < 0.01, cluster corrected | | | | | | |
|---|---|---|---|---|---|---|
| Region | Hemisphere | Size (mm$^3$) | Peak X | Peak Y | Peak Z | t |
| Middle Occipital Gyrus | Left | 2960 | -52 | -73 | -5 | 5.56 |
| Cuneus | Medial | 2803 | 0 | -91 | 29 | 4.85 |

| Correlation between Counting List & Number Preference: r(14) > 0.62, p < 0.01, cluster corrected | | | | | | |
|---|---|---|---|---|---|---|
| Region | Hemisphere | Size (mm$^3$) | Peak X | Peak Y | Peak Z | r |
| Intraparietal Sulcus | Left | 1925 | -45 | -43 | 49 | 0.78 |
| Superior Parietal Lobule | Left | 3411 | -27 | -67 | 52 | 0.81 |
| Superior Temporal Gyrus | Right | 2405 | 45 | 14 | -23 | 0.80 |
| Caudate | Right | 1900 | 9 | 17 | 7 | 0.81 |

**Table S3.** Sequence Processing Results. Note: for the Alphabet Sequences > Counting Sequences blocked contrast, a cluster that spanned the precuneus and angular gyrus has two peaks listed, one for the precuneus and one for the angular gyrus. The angular gyrus has been implicated in numerical processing, reading, and ordinal information so the peak coordinates for that area within the larger cluster are listed separately to better aid future research. The accompanying hemisphere and size values correspond to the entire cluster.

| **Counting (Parametric Effect): t(42) > 2.70, p < 0.01, cluster corrected** | | | | | | |
|---|---|---|---|---|---|---|
| Region | Hemisphere | Size ($mm^3$) | Peak X | Peak Y | Peak Z | t |
| Intraparietal Sulcus | Right | 10940 | 48 | -40 | 52 | 5.48 |
| Intraparietal Sulcus | Left | 4891 | -45 | -46 | 46 | 4.99 |
| Inferior Frontal Gyrus | Left | 5140 | -48 | 3 | 3 | 5.34 |
| Anterior Temporal Lobe/Superior Temporal Gyrus | Right | 8724 | 57 | -19 | 1 | 6.36 |
| Anterior Temporal Lobe/Superior Temporal Gyrus | Left | 16433 | -54 | -7 | 1 | 7.53 |
| Medial Frontal Gyrus | Medial | 3543 | -3 | -7 | 58 | 5.14 |
| Precentral Gyrus | Left | 6234 | -48 | -13 | 40 | 4.98 |
| Middle Frontal Gyrus | Left | 4517 | -39 | 32 | 28 | 4.43 |
| Putamen | Medial | 6971 | -18 | -1 | 13 | 4.66 |

| **Alphabet (Parametric Effect): t(42) > 2.70, p < 0.01, cluster corrected** | | | | | | |
|---|---|---|---|---|---|---|
| Region | Hemisphere | Size ($mm^3$) | Peak X | Peak Y | Peak Z | t |
| Inferior Parietal Lobule | Right | 2889 | 49 | -64 | 40 | 3.60 |
| Inferior Frontal Gyrus | Right | 1356 | 58 | 11 | 25 | 3.77 |
| Inferior Frontal Gyrus/Insula | Right | 4014 | 30 | 23 | 7 | 4.26 |
| Inferior Frontal Gyrus/Caudate | Left | 6055 | -15 | -10 | 19 | 4.44 |
| Caudate | Right | 2672 | 12 | 14 | 7 | 4.46 |
| Caudate | Left | 5302 | -12 | 14 | 4 | 4.51 |
| Middle Frontal Gyrus | Right | 6298 | 33 | 20 | 46 | 4.31 |
| Middle Frontal Gyrus | Right | 1769 | 21 | 14 | 61 | 3.81 |

| | | | | | | |
|---|---|---|---|---|---|---|
| Superior Frontal Gyrus | Right | 3206 | 24 | 38 | 28 | 4.39 |
| Precentral Gyrus | Left | 3047 | -48 | 11 | 4 | 4.03 |
| Medial Frontal Gyrus | Left | 677 | -18 | 35 | 28 | 3.25 |
| Cingulate Gyrus | Right | 30363 | 3 | 29 | 31 | 4.83 |
| Cingulate Gyrus | Medial | 6712 | 0 | -16 | 25 | 3.96 |
| Anterior Temporal Lobe/Superior Temporal Gyrus | Right | 2401 | 60 | -4 | 7 | 4.49 |
| Anterior Temporal Lobe/Superior Temporal Gyrus | Right | 1900 | 51 | 8 | -2 | 4.12 |
| Anterior Temporal Lobe/Superior Temporal Gyrus | Left | 1421 | -54 | -13 | 4 | 4.00 |
| Precuneus | Medial | 2401 | 0 | -73 | 43 | 3.91 |
| Occipital Cortex | Right | 17116 | 5 | -82 | 16 | 5.77 |
| Occipital Cortex | Right | 16173 | 9 | -69 | 16 | 5.52 |
| Occipital Cortex | Medial | 14327 | 0 | -76 | 10 | 5.94 |
| Occipital Cortex | Left | 11260 | -12 | -70 | 10 | 5.29 |
| Occipital Cortex | Left | 10869 | -3 | -79 | 19 | 7.01 |
| Occipital Cortex | Right | 6528 | 15 | -73 | -17 | 5.14 |
| Cerebellum/Ventral Temporal Cortex | Right | 9148 | 15 | -43 | -18 | 5.26 |
| Cerebellum/Ventral Temporal Cortex | Left | 8805 | -6 | -61 | -12 | 5.16 |
| Cerebellum/Ventral Temporal Cortex | Left | 1673 | -30 | -58 | -27 | 3.87 |
| Cerebellum | Right | 9404 | 9 | -48 | -1 | 4.70 |
| Thalamus | Medial | 4573 | 0 | -7 | 16 | 4.24 |
| Thalamus | Right | 4638 | 12 | -13 | 16 | 4.98 |
| Thalamus | Right | 1656 | 15 | -22 | 16 | 4.34 |
| Lentiform Nucleus | Right | 990 | 27 | -19 | -2 | 3.65 |
| Claustrum | Left | 1999 | -27 | 23 | 1 | 4.28 |

| Alphabet Sequences > Counting Sequences: t(42) > 2.70, p < 0.01, cluster corrected | | | | | | | |
|---|---|---|---|---|---|---|---|
| Region | | Hemisphere | Size (mm$^3$) | Peak X | Peak Y | Peak Z | t |
| Precuneus/Angular Gyrus: | Precuneus Peak | Left | 3515 | -18 | -73 | 22 | 4.40 |
| | Angular Gyrus Peak | | | -36 | -58 | 37 | 3.40 |
| Inferior Frontal Gyrus | | Left | 1119 | -48 | 5 | 28 | 3.79 |
| Middle Temporal Gyrus | | Left | 1511 | -54 | -34 | 4 | 4.50 |
| Cuneus | | Right | 1574 | 27 | -76 | 10 | 4.03 |

**Table S4.** Correlations Between Neural Maturity and Number Skills

| **Neural Maturity/Number Skills Correlation: r(37) > 0.41, p < 0.01, cluster corrected** | | | | | | |
|---|---|---|---|---|---|---|
| Region | Hemisphere | Size ($mm^3$) | Peak X | Peak Y | Peak Z | r |
| Intraparietal Sulcus | Left | 2311 | -33 | -55 | 49 | 0.64 |
| Precuneus | Right | 699 | 15 | -64 | 49 | 0.48 |
| Precuneus | Left | 1064 | -12 | -58 | 55 | 0.58 |

| **Neural Maturity/Number Skills Correlation (Controlling Letter Skills) : r(37) > 0.41, p < 0.01, cluster corrected** | | | | | | |
|---|---|---|---|---|---|---|
| Region | Hemisphere | Size ($mm^3$) | Peak X | Peak Y | Peak Z | r |
| Intraparietal Sulcus | Left | 759 | -36 | -40 | 40 | 0.53 |
| Medial Frontal Gyrus | Left | 906 | -9 | 59 | 13 | 0.56 |
| Posterior Middle Temporal Gyrus | Left | 1232 | -45 | -73 | 16 | 0.58 |
| Precuneus | Medial | 675 | 0 | -73 | 43 | 0.55 |
| Occipital Cortex | Right | 930 | 54 | -46 | -5 | 0.63 |
| Limbic Lobe | Left | 608 | -27 | 2 | -38 | 0.55 |

**Table S5.** Results for Comparison of Known vs Unknown Number Words

| Known Numbers > Unknown Numbers: t(33) > 2.73, p < 0.01, cluster corrected | | | | | | |
|---|---|---|---|---|---|---|
| Region | Hemisphere | Size ($mm^3$) | Peak X | Peak Y | Peak Z | t |
| Hippocampus/Thalamus | Right | 3818 | 15 | -28 | 1 | 4.32 |
| Ventral Temporal Cortex | Right | 5461 | 33 | -43 | -11 | 4.99 |
| Occipital Cortex | Right | 1849 | 9 | -85 | -8 | 3.65 |

| Unknown Numbers > Known Numbers: t(33) > 2.73, p < 0.01, cluster corrected | | | | | | |
|---|---|---|---|---|---|---|
| Region | Hemisphere | Size ($mm^3$) | Peak X | Peak Y | Peak Z | t |
| Intraparietal Sulcus | Right | 2224 | 42 | -52 | 52 | 4.22 |
| Intraparietal Sulcus | Left | 4715 | -39 | -58 | 37 | 4.34 |
| Middle Frontal Gyrus | Left | 2290 | -36 | 35 | 22 | 4.33 |
| Precentral Gyrus | Left | 3961 | -45 | 8 | 10 | 5.19 |
| Anterior Cingulate Cortex | Left | 3993 | -3 | 23 | 46 | 4.85 |
| Cingulate Gyrus | Right | 4566 | 6 | -22 | 28 | 4.62 |
| Anterior Temporal Lobe/Superior Temporal Gyrus | Left | 7598 | -54 | -10 | 1 | 6.49 |
| Precuneus | Left | 2910 | -12 | -67 | 37 | 4.39 |
| Precuneus | Right | 3088 | 12 | -61 | 31 | 4.95 |

**Table S6**. Numerosity Task Results.

| Hard Ratio + Easy Ratio across Children & Adults: t(64) > 2.66, p < 0.01, cluster corrected | | | | | | |
|---|---|---|---|---|---|---|
| Region | Hemisphere | Size ($mm^3$) | Peak X | Peak Y | Peak Z | t |
| Inferior Parietal Cortex | Right | 9213 | 24 | -64 | 28 | 4.89 |
| Inferior Parietal Cortex | Left | 10255 | -21 | -67 | 37 | 5.64 |
| Inferior Frontal Gyrus/Insula | Right | 4678 | 33 | 17 | 7 | 4.24 |
| Inferior Frontal Gyrus/Insula | Left | 4947 | -27 | 17 | 10 | 4.78 |
| Precentral Gyrus | Right | 3827 | 39 | -1 | 31 | 4.54 |
| Precentral Gyrus | Left | 5974 | -36 | -16 | 58 | 4.50 |
| Precentral Gyrus | Left | 5791 | -42 | -1 | 31 | 5.18 |
| Anterior Cingulate Cortex | Medial | 25957 | -6 | 5 | 43 | 5.12 |
| Occipital Cortex | Bilateral | 121035 | -9 | -82 | -11 | 7.21 |
| Thalamus | Medial | 22259 | -21 | -28 | -2 | 6.27 |

| Hard Ratio > Easy Ratio across Children & Adults: t(64) > 2.66, p < 0.01, cluster corrected | | | | | | |
|---|---|---|---|---|---|---|
| Region | Hemisphere | Size ($mm^3$) | Peak X | Peak Y | Peak Z | t |
| Intraparietal Sulcus | Right | 19277 | 30 | -52 | 37 | 7.28 |
| Intraparietal Sulcus | Left | 7339 | -45 | -43 | 34 | 5.45 |
| Inferior Frontal Gyrus/Insula | Left | 11871 | -33 | 14 | 7 | 10.67 |
| Inferior Frontal Gyrus/Insula | Right | 17523 | 30 | 17 | 4 | 11.76 |
| Precentral Gyrus | Right | 13610 | 42 | -1 | 31 | 6.97 |
| Middle Frontal Gyrus | Right | 5345 | 27 | -4 | 58 | 4.90 |
| Anterior Cingulate Cortex | Medial | 31604 | 3 | 8 | 46 | 9.97 |
| Thalamus | Medial | 3878 | 12 | -7 | 7 | 5.01 |
| Occipital Cortex | Right | 87624 | 15 | -85 | 1 | 11.54 |

**Supplement 4**

Supplement 4 depicts additional results from the Word Processing Task. Details of these regions can be found in Supplement 3, Table S2.

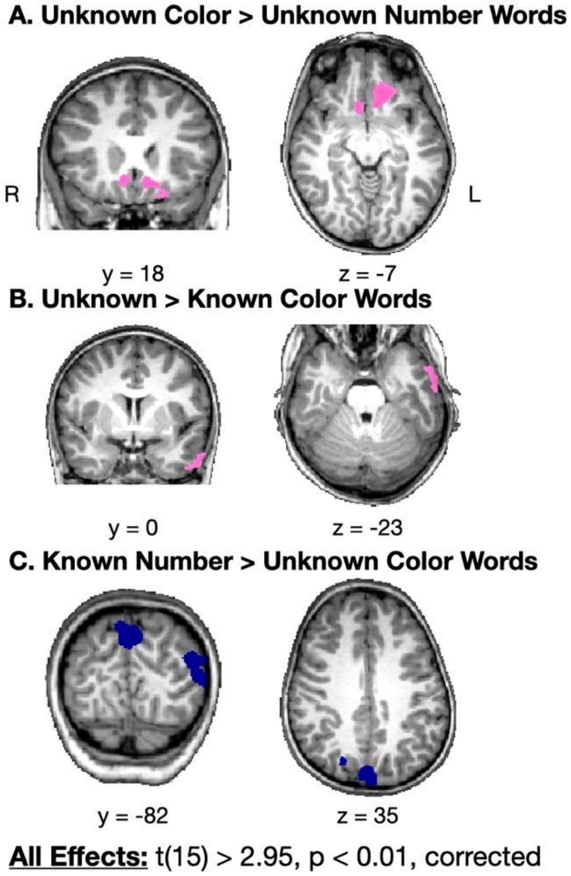


**Figure S1.** Regions that showed significant effects in the follow-up whole-brain contrasts for the word processing task. (A) Unknown Color > Unknown Number Words. (B) Unknown > Known Color Words. (C) Known Number > Unknown Color Words. The reverse contrasts did not show any significant effects, and all significant effects are visible in the images shown here. Details of these regions can be found in Supplement 3, Table S2.

**Supplement 5**

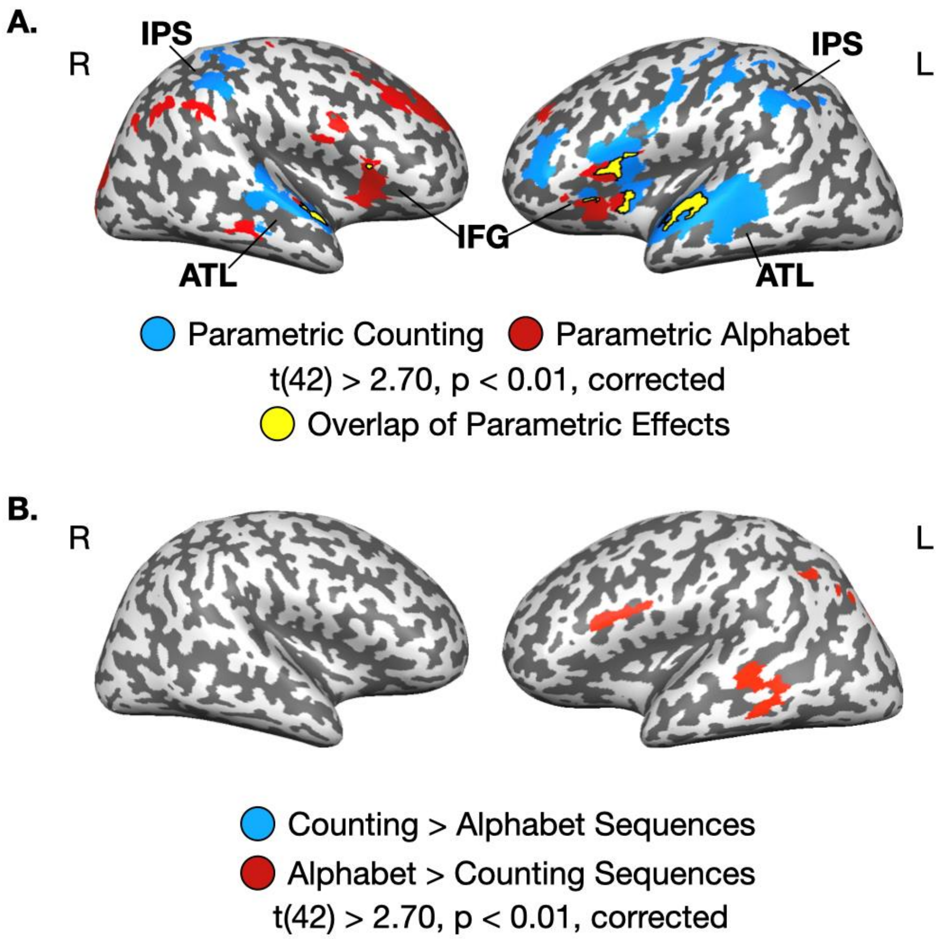


**Figure S2.** Children's neural responses while listening to counting sequences (blue) and alphabet sequences (red). (A) Regions that show an increase in activation as children listen to the progression of sequences (parametric effects of sequence processing). (B) Regions that show a difference in overall activation between listening to counting sequences vs alphabet sequences (counting sequences vs alphabet sequences). A full list of all coordinates and regions for both panels is presented in Supplement 3, Table S3. All results are shown at a threshold of $p < 0.01$, corrected, with an applied cluster threshold of 25 $mm^3$.

**Supplement 6**

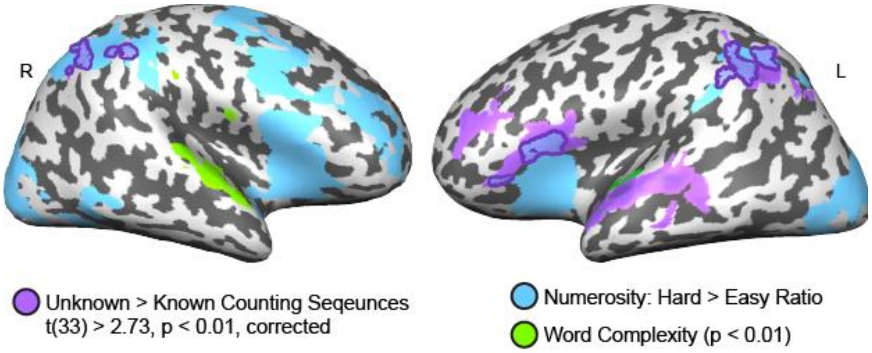


**Figure S3.** Effect of word complexity (green) compared to numerosity processing regions (blue) and regions that showed greater activation for Unknown counting sequences (purple)

Regions that were sensitive to word complexity (green) were identified by testing for a parametric effect of the number of syllables in the stimuli presented in the word processing task and the natural counting task. Importantly the regions that showed an effect of word complexity did not overlap with those that showed a novelty effect for number words (Unknown > Known counting sequences), suggesting that the novelty effect is not confounded by greater complexity of the larger, Unknown number words.

**Supplement 7**

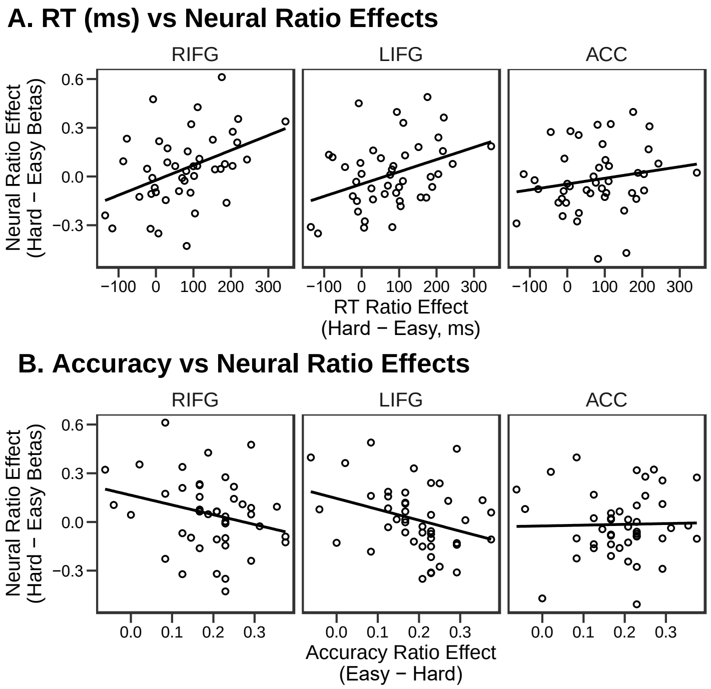


**Figure S4.** Correlations between children's neural ratio effects in the numerosity network and their performance on the numerosity comparison task. Behavioral performance is assessed through (A) response time ratio effects and (B) accuracy ratio effects. Comparable figures for the IPS are presented in the main text, Figure 5B.

Beta values for the easy and difficult comparison conditions were extracted from spheres surrounding the peak coordinates in the left and right IFG and the ACC from the children's numerosity discrimination data (numerosity discrimination task > baseline). The ratio effect in children's response time (RT; RT for difficult trials – RT for easy trials) was correlated with children's neural ratio effect (beta for difficult ratios – beta for easy ratios) in the left and the right IFG, but not in the ACC (Figure S1A; left IFG: r(42) = 0.38, t(42) = 2.63, p = 0.012; right IFG: r(42) 0.42, t(43) = 3.02, p = 0.004; ACC: r(42) = 0.18, t(42) = 1.20, p = 0.239). This suggests that children who showed a larger difference in RT between ratios showed a larger neural ratio effect in bilateral IFG. The ratio effect in children's accuracy (accuracy for easy trials – accuracy for difficult trials) was correlated with children's neural ratio effect in the left IFG, was moderately correlated with children's neural ratio effect in the right IFG, and was not corrected in ACC (Figure S1B; left IFG: r(43) = -0.32, t(43) = -2.20, p = 0.033; right IFG: r(43) = -0.26, t(43) = -1.80, p = 0.079; ACC: r(43) = 0.02, t(43) = 0.15, p = 0.878). Children who showed a larger difference in accuracy between ratios showed a smaller neural ratio effect in the left IPS. This shows that behavioral performance on the numerosity discrimination task relates to neural activity in the bilateral IFG.

**Supplement 8**

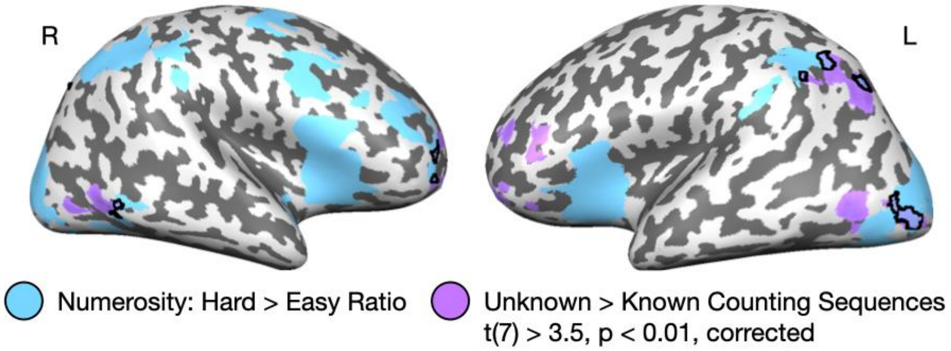


**Figure S5.** Comparison of the main effect of numerosity comparison ratio and unknown > known counting sequences in children who had not yet acquired the counting principle (“subset knowers”). A comparable figure for the whole group is shown in Figure 6B.

Knowledge of the cardinality principle was assessed using the Give-N task. Eight children did not demonstrate knowledge of the cardinality principle so were classified as subset knowers. A comparison of Unknown to Known counting sequences in this set of children revealed sensitivity for Unknown counting sequences in the left IPS. This region is similar to that identified in the full group of children and shows functional overlap with numerosity processing, suggesting that the left IPS is involved in the acquisition of number words prior to the acquisition of the cardinality principle.

**Supplement 9**

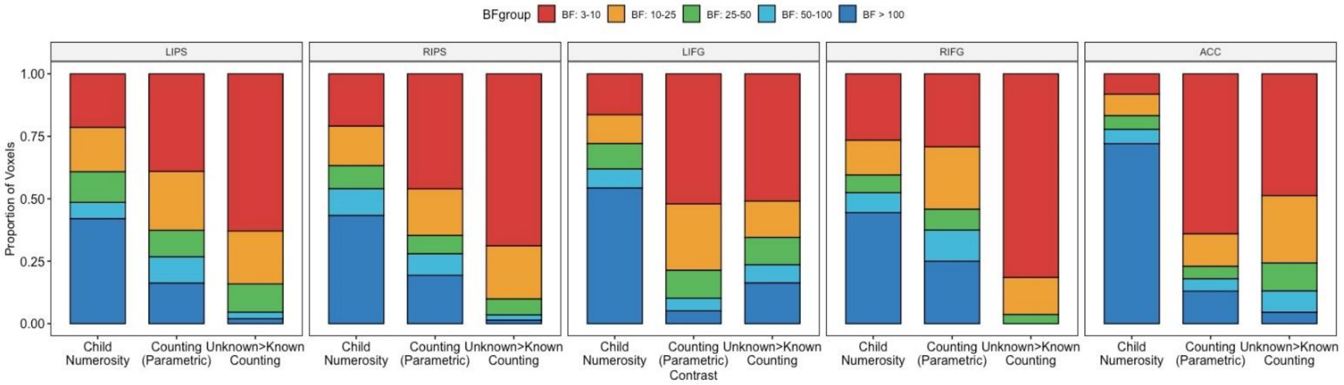


**Figure S6.** Distribution of $BF_{10}$ values in voxels that showed substantial effect of both adult numerosity processing and the child numerosity and count processing effects.

Within voxels that showed substantial evidence of both adult numerosity effects and child numerosity and counting effects ($BF_{10} \geq 3$), we examined the proportion of voxels that fell into 5 ranges of $BF_{10}$ values: 3-10, 10-25, 25-50, 50-100, and >100 (Figure S3). Stronger $BF_{10}$ values were seen for the child numerosity effect than the counting effects. This is unsurprising given that the child numerosity effect and the adult numerosity ratio effect draw on data from the same task (but note that here we used independent groups). In contrast, the counting data come from a completely different task and targeting different numerical representations (spoken number words, counting). 54% - 61% of IPS voxels showed a $BF_{10} \geq 10$ for the parametric counting effect (LIPS: 61%, RIPS: 54%, LIFG: 48%, RIFG: 71%, ACC: 36%) and 31% - 37% of IPS voxels showed a $BF_{10} \geq 10$ for unknown count words (LIPS: 37%, RIPS: 31%, LIFG: 49%, RIFG: 19%, ACC: 51%). In sum, of the voxels that showed evidence of both adult numerosity processing and early childhood sequence processing ($BF_{10} \geq 3$ for both effects), at least 19% of voxels in the IFG and at least 31% of voxels in the IPS showed strong evidence of count sequences processing ($BF_{10} \geq 10$). In other words, in voxels where there are effects of both numerosity processing and early childhood number word representations, the evidence for each effect is robust.
